# MetaGeneMark-2: Improved Gene Prediction in Metagenomes

**DOI:** 10.1101/2022.07.25.500264

**Authors:** Karl Gemayel, Alexandre Lomsadze, Mark Borodovsky

**Affiliations:** School of Computational Science and Engineering, Georgia Tech, Atlanta, GA 30332, USA; Wallace H Coulter Department of Biomedical Engineering, Georgia Tech and Emory University, Atlanta, GA 30332, USA

**Keywords:** metagenome sequence analysis, gene prediction, regulatory motif, model merging, stop codon reassignment

## Abstract

Accurate prediction of protein-coding genes in metagenomic contigs presents a well-known challenge. Particularly difficult is to identify short and incomplete genes as well as positions of translation initiation sites. It is frequently assumed that initiation of translation in prokaryotes is controlled by a ribosome binding site (RBS), a sequence with the Shine-Dalgarno (SD) consensus situated in the 5’ UTR. However, ∼30% of the 5,007 genomes, representing the RefSeq collection of prokaryotic genomes, have either non-SD RBS sequences or no RBS site due to physical absence of the 5’ UTR (the case of leaderless transcription). Predictions of the gene 3’ ends are much more accurate; still, errors could occur due to the use of incorrect genetic code. Hence, an effective gene finding algorithm would identify true genetic code in a process of the sequence analysis. In this work prediction of gene starts was improved by inferring the GC content dependent generating functions for RBS sequences as well as for promoter sequences involved in leaderless transcription. An additional feature of the algorithm was the ability to identify alternative genetic code defined by a reassignment of the TGA stop codon (the only stop codon reassignment type known in prokaryotes). It was demonstrated that MetaGeneMark-2 made more accurate gene predictions in metagenomic sequences than several existing state-of-the-art tools.

## Introduction

A 2007 study estimated that less than 1% of bacteria inhabiting soil or aquatic environments could be cultured in a lab (National Research Council 2007). Nonetheless, the shotgun sequencing of DNA of microbial communities was introduced with the idea to get insights into biology and ecology of vast majority of prokaryotic species through decoding of the metagenomic fragments. Accurate gene prediction in anonymous metagenomic sequences is facing a challenge. The sequences are lacking a context of a much longer genomic sequence, that would be used for estimation of species-specific parameters of the algorithm. Therefore, *ab initio* gene prediction in metagenomic fragments requires special techniques for construction of the models of protein-coding regions (Besemer and Borodovsky 1999; Noguchi et al. 2006; Hoff et al. 2009; Rho et al. 2010; Zhu et al. 2010; Kelley et al. 2012). Additional efforts are needed to improve gene start prediction as well as to identify possible stop codon reassignments (Noguchi et al. 2008; Hu et al. 2009; Hyatt et al. 2012). Here, following the earlier publication of MetaGeneMark (Zhu et al. 2010), we present a new algorithm and software tool, MetaGeneMark-2. The algorithm design has taken into account introduction of multiple models of RBS and promoter sites made upon development of GeneMarkS-2 (Lomsadze et al. 2018). The new algorithm is using the G+C composition and taxonomy dependent models of sequences surrounding gene starts. It also uses locally adjusted G+C content dependent models for detecting short and incomplete genes. Finally, it is aware of a possible stop codon reassignment in a contig of unknown nature and attempts to detect possible reassignment during sequence analysis. We show that the new tool improves gene start prediction accuracy in comparison with earlier developed metagenomic gene finders.

## Materials

### Data sets for derivation of the MetaGeneMark-2 parameter generating functions

We used a set of 5,491 representative RefSeq genomes (5,238 bacteria and 253 archaea) selected by NCBI. In this set, all archaeal and 5,137 bacterial genomes had the ‘standard’ genetic code (code 11); TGA was coding for tryptophan (Trp) amino acid (genetic code 4) in 101 bacterial genomes.

### Sets of genes with experimentally verified starts

Positions of translation starts were experimentally verified by N-terminal protein sequencing in several species. In our tests we used sets of verified genes from the following species of bacteria: *Synechocystis sp*. (Sazuka et al. 1999), *E. coli* (Rudd 2000; Zhou and Rudd 2013), *M. tuberculosis* (Lew et al. 2011), *R. denitrificans* (Bland et al. 2014) and *D. deserti* (de Groot et al. 2014), as well as the species of archaea *A. pernix* (Yamazaki et al. 2006), *H. salinarum* and *N. pharaonis* (Aivaliotis et al. 2007).

### Sets of genes verified by the StartLink+ predictions

The sets of genes with experimentally verified starts are not large due to the costs of their production. We needed additional tests sets due to the variety of patterns existing in the gene upstream regions. In the *Enterobacterales* clade with predominantly mid-GC genomes, the RBS sites have the Shine-Dalgarno consensus. In the bacteria of the *FCB group* the RBS sequences have AT-rich ‘non-canonical’ consensus (Lomsadze et al. 2018). In archaeal genomes large number of genes are expressed in leaderless transcripts (with no upstream RBS sites). Also, leaderless transcription is ubiquitous in *Actinobacteria* with high-GC genomes. To get additional test sets we used the following approach. In our previous work (Gemayel et al. 2021) we have shown that predictions of gene starts made by StartLink+ have low error rates, ∼1%. Therefore, we used StartLink+ to make computational predictions of gene starts in 427 genomes including 108 archaeal, and 319 bacterial ones (104 *Enterobacterales*, 111 *Actinobacteria*, and 104 *FCB group*). We used the sets of genes predicted by StartLink+ as additional test sets.

### Set of genomes for testing accuracy of identification of the role of TGA codon

We used 11,688 RefSeq genomes and annotation of 405 archaea and 11,265 bacteria (www.ncbi.nlm.nih.gov/genome/browse#!/prokaryotes/). All the genomes but 167 bacterial genomes with genetic code ‘4’ had the standard genetic code (code 11). To increase number of genomes with code ‘4’ with TGA reassigned to Trp we added 18 GenBank genomes of the *Hodgkinia* genus.

## Methods

### Modeling of the protein coding regions

To infer functions generating parameters of the GC-dependent models for protein coding regions MetaGeneMark-2 uses the same approach as in MetaGeneMark (Zhu et al. 2010). As a novel addition, the GC-dependent parameter generating functions were inferred for gene upstream regulatory sequences (RBS & promoter sites), for spacer length distributions, for frequency distributions of start codons, and for sequences of the gene start contexts. The model structures were selected in agreement with the genome’s phylogenetic classification, i.e., groups A, B, C, D, and X, introduced in GeneMarkS-2 (Lomsadze et al. 2018). To explicitly indicate archaeal genomes, we slightly modified this nomenclature to feature larger number of groups Γ ∈ {*A, A*^*∗*^, *B, C, D*^*∗*^, *X*}, where the asterisks indicate archaeal groups.

To construct the generating functions for parameters of regulatory signals we used sets of parameters of the models derived by GeneMarkS-2 for genomes of isolates of prokaryotic species. Let 𝒜 and ℬ be the sets of archaeal and bacterial genomes, respectively. For each genome in 𝒜 and ℬ we built a genome-specific GeneMarkS-2 model indexed by the group label Γ. In addition, we constructed models of sequences situated near gene starts, as described below, for each GC bin *b* = [*b*_*l*_, *b*_*u*_), defined by its lower and upper limits *b*_*l*_ and *b*_*u*_ (e.g., *b* = [30,31) or *b* = [40,45)). For derivation of generating functions for parameters of the RBS and promoter motif models we used GC bins with width 5%, while the bin width of 1%, i.e. (30, 31), (31, 32), …, (69, 70) was used in derivation of all the other generating functions.

### Modeling of the start codons frequency and the gene start context

Let *p*_*s*_, where *s* ∈ {ATG, GTG, TTG}, be the start codon frequencies observed in genes predicted by GeneMarkS-2 in each genome. In the 𝒜 and ℬ genomes we observed that the values of {*p*_*s*_} changed across the entire GC range showing group-specific patters (Fig. 1, top panel).

**Figure 1.**
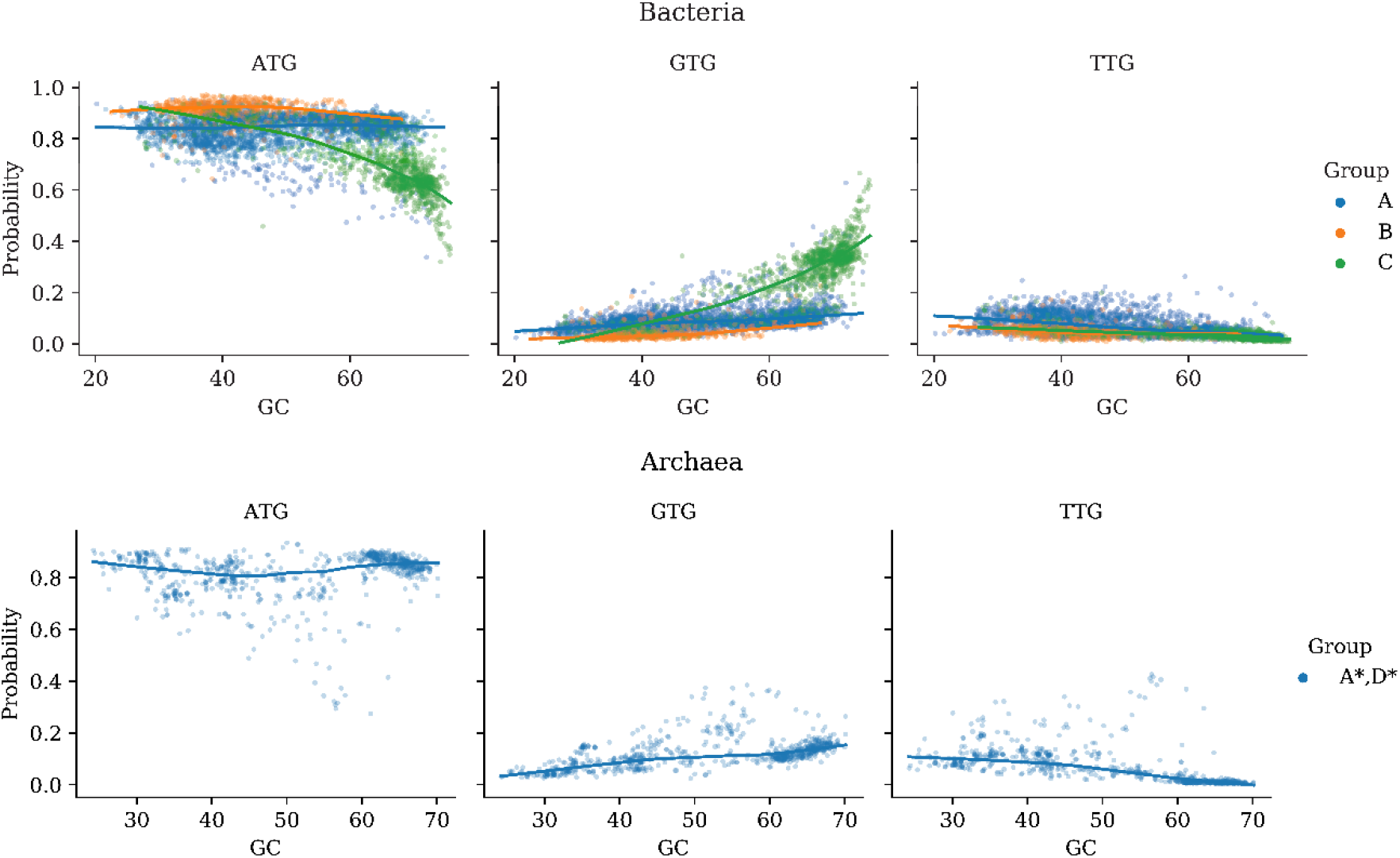
The probabilities of the three start codons as functions of GC content estimated from the sets of genes predicted by GeneMarkS-2 in 5,491 archaeal and bacterial genomes. The data for small size archaeal groups A* and D* (bottom panel) were combined together.

To construct generating functions for probabilities of *start codons* and probabilities of nucleotides in the *gene start context* as a function of the sequence GC content in each genome group Γ we used LOWESS, the locally weighted scatterplot smoothing (Cleveland 1981). Each regression curve was “discretized” by computing average frequencies of ATG, GTG or TTG per a GC bin *b*. Formally, the probability of a codon *s* for a group Γ genomes in a GC bin *b* was estimated as

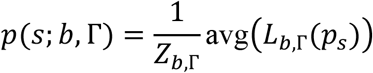

where *L*_*b*,Γ_(*p*_*s*_) was determined by the LOWESS smoothing for the *p*_*s*_ values from group Γ in the GC bin *b*, the *avg* function computed mean values, and *Z*_*b*,Γ_ was a normalization constant. A 15 nt sequence occupying positions -3 to 12, where the first nucleotide of a start codon was situated in position 1, was defined as the gene start-context sequence. Parameters of a model for the start-context sequence, the second order positional Markov chain, were produced by corresponding generating functions. To derive these functions for each group of genomes Γ we approximated positional frequencies of 64 nucleotide triplets determined in GC dependent bins by the LOWESS smoothing.

### Positional frequency models of RBS, promoter sites and spacer length distributions

Parameters of the RBS and promoter site models were estimated by computing positional frequencies of nucleotides in alignments of predicted regulatory sequences. These parameters could vary even among species *that belong to the same taxonomic group and have the same GC content*. The observed variations in frequency matrices were of several kinds (1) minor fluctuations in frequencies without change of the consensus in each position; (2) moderate frequency fluctuations that would cause a change of the consensus in one or two positions; (3) fluctuations of frequencies that would make the motif consensus in one genome appear to be shifted relatively to the motif consensus in another genome, e.g., AGGAGG vs AAGGAG. Note that in a model for a group of genomes a length of the merged motif model should be sufficient to comprise all the motif models derived for particular genomes. Also, we should keep in mind that shorter consensus could have multiple options to appear in a longer consensus, e.g., the 4 position consensus GGAG could appear in a 6 position consensus as XXGGAG, XGGAGX, or GGAGXX, where X designates *any* type of nucleotide.

### Merging site models within the motif clusters

Suppose we have N_A_ genomes in a taxonomic group A; let assume also that all N_A_ genomes belong to the same GC content bin. We reduce the number of the models by introducing merged models for genomes whose models have similar consensus sequences. Then, we look for a minimal number of genome clusters, *K*, satisfying the condition that within each cluster, *h =* 0,1,2, … *K* any pair of the motif consensus sequences would differ at most in αtwo positions. For instance, let have four genomes, N_A_ = 4, whose RBS models have the following consensus sequences:

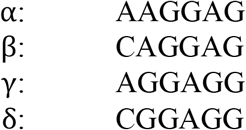

Here genomes *α* and *β* belong to one cluster, and *γ* and *δ* -belong to another one. Parameters of one merged RBS model are estimated from all the predicted RBS sequences in genomes *α* and *β*, while parameters of another merged model are estimated from the RBS sequences predicted in genomes *γ* and *δ*. The spacer length distributions are derived by collecting the lengths of predicted spacers in each pair of the genomes.

Formally, let *∧* be the entire set of RBS models derived by running GeneMarkS-2 on a set of genomes. Let variable *θ* = (*b*, Γ) represent a specific GC bin *b* and genome group Г. For each *θ*, we will have a set of merged models described by the following components: i/ prior probabilities of each motif cluster in *θ, p*(*h; θ*), *h =* 0,1,2, … m_θ_; ii/ merged *motif models M*_*θ,h*_ for each cluster *h*, and iii/ merged spacer length distributions *S*_*θ,h*_ for each cluster *h*. Similar definitions are used to for merged promoter site models.

Therefore, for an anonymous metagenomic contig that falls within a given GC bin *b* we have a rather small number of the GC specific *merged* motif/spacer models defined for each genome group Г. In Fig. 2 we show an example of a merged RBS model for the group A bacterial genomes with GC bin *b* = [35,40), i.e., *θ* = ([35,40), *A*). The top two panels show merged motif models for clusters h=0 and h=1. In each panel, the first plot shows relative entropy logo of the merged motif model. The next four plots show distributions of positional frequencies of each nucleotide. The heights of the boxes show the standard deviations among the nucleotide frequencies within the merged model.

**Figure 2.**
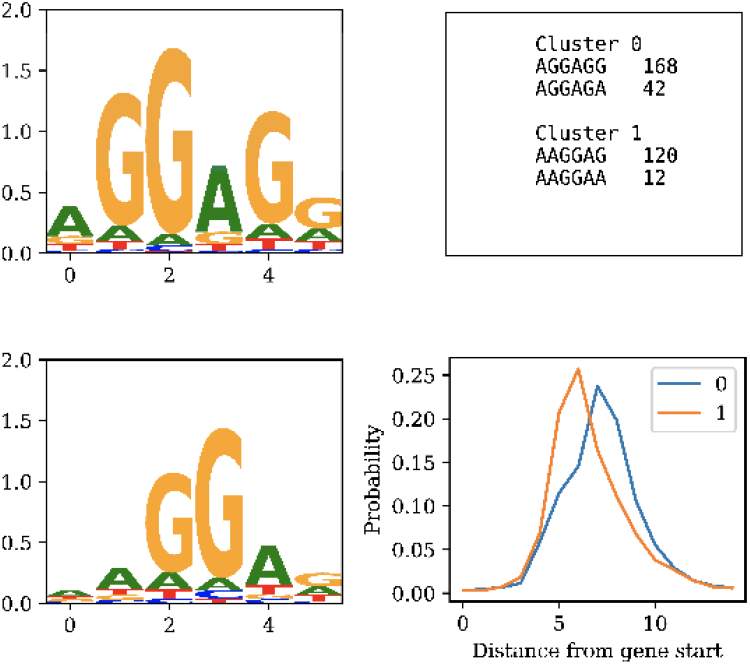
The two-clustered Shine-Dalgarno RBS model made in the GC content range [45, 50] (see text for details)

In the bottom panel the first plot shows consensus sequences of the models related to clusters h=0 and h=1, with the numbers of merged models per each consensus. The second plot shows prior probabilities of each merged model. The last plot depicts the merged spacer length distributions for each merged model.

An example of a merged bacterial promoter model for *θ* = ([60,65), *C*) is shown in Fig. 3. The TANNNT promoter consensus is typical for the GeneMarkS-2 models identified in the group C genomes (Lomsadze et al. 2018); this type of pattern was also observed experimentally in the group C genome *M. tuberculosis* (Cortes et al. 2013).

**Figure 3.**
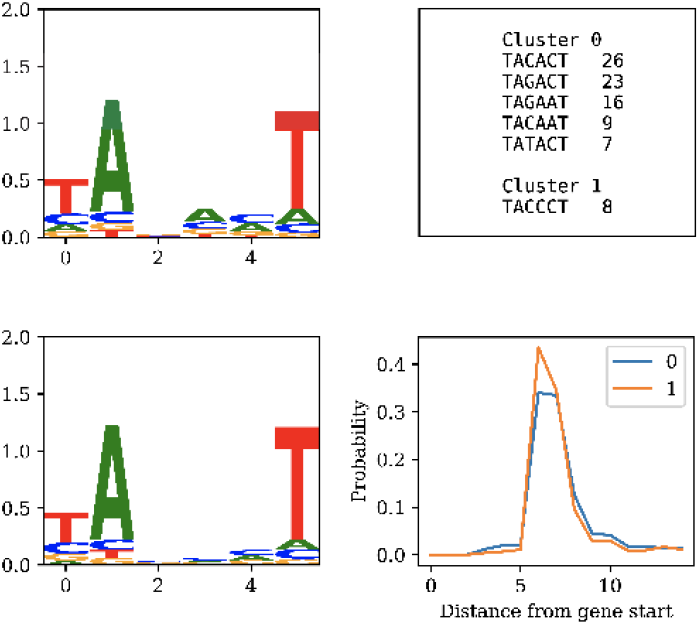
The two-clustered bacterial promoter model made in the GC content range [60, 65] (see text for details).

### Gene prediction

The task of gene prediction in prokaryotic genomic sequence was formalized with application of the theory of hidden Markov models (HMMs). The simplifying premise of this modeling theory is that the genomic sequence is emitted, segment by segment, from a sequence of hidden states corresponding to protein-coding and non-coding regions. Therefore, the task was reformulated as finding a sequence of hidden states of a generalized HMM and their durations (the lengths of the emitted segments) which joint probability to occur along with the whole emitted genomic sequence would be maximal in comparison with any other sequence of hidden states.

Such a task was solved by the GeneMark.hmm and the self-training GeneMarkS algorithms (Lukashin and Borodovsky 1998; Besemer et al. 2001) with implementation of the Viterbi algorithm for the semi-Markov HMM. GeneMarkS-2, also designed for isolated genomes, introduced differences with the earlier algorithms (see Fig. 1 in Lomsadze et al. 2018). First, the generating functions delivered initial parameters of the models of protein-coding regions based on local GC content (an ORF GC content), second, multiple models of regulatory sequences (RBS and promoters) could be used simultaneously, and, third, a model was introduced for a sequence (15 nt) downstream of a putative start codon (a downstream signature).

Each ORF sequence longer than 90 nt was assigned to a GC bin *b*_*c*_ based on its GC content. For each candidate start codon, the upstream nucleotide sequence *U* with length, *L*, (20nt for bacteria and 40 nt for archaea) was determined.

Here we modified definition of taxonomic groups introduced in the GeneMarkS-2 publication (Lomsadze et al, 2018) by separating bacteria and archaea, as well as by making distinction between groups having genetic code 11 and 4. For instance, collection *G*_11_ = {*A, A*^*∗*^, *B, C, D*^*∗*^, *X*}_11_, where asterisks indicate archaea, and collection *G*_4_ = {*A, C*}_4_. In sections 3.2 and 3.3 we described how to determine parameters for GC-dependent models for each group in the collections *G*_11_and *G*_4_.

To compute a probability of a metagenomic contig, we assumed that the contig belonged to a genome from group Γ, and used i/ GC content specific models of the protein-coding region for each ORF and ii/ several cluster specific models of upstream regions that were 21nt long for bacteria and 40 nt long for archaea. In comparison with GeneMarkS-2 there was an additional step of optimization. For each contig the task was to select a merged model of the regulatory region delivering the highest joint probability of the sequence of hidden states and the whole contig sequence.

In each MetaGeneMark-2 run we assumed that a metagenomic sequence belongs to one of the groups Γ from collections *G*_11_or *G*_4_. We calculated probability of generating observed sequence by group specific models for all possible Γ and the prediction with highest probability value was selected as the final prediction in this sequence:

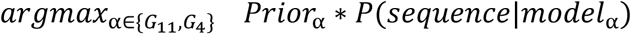

where *Prior*_*a*_ was prior probability for groups from *G*_11_and *G*_4_.

The Viterbi algorithm was implemented with the two main differences: i/ the parameters of the models of RBS, promoter, extended (group X) upstream signature, start codon and downstream signature were determined by generating functions dependent on GC content of each open reading frame; ii/ the GC content dependent models of RBS and promoters could be represented by several sub-models (created by merging sets of single motifs into motif clusters, see below).

Implementation of the first feature was akin to derivation of GC dependent models of the protein-coding regions in GeneMarkS-2. The second feature was implemented within the generalized Viterbi algorithm finding the optimal parse of a contig into coding and non-coding regions. The algorithm had to consider all possible combinations of gene start candidates in all possible chains of open reading frames (ORFs) existed in the contig. The dynamic programming problem has essentially quadratic complexity even if several alternative starts are considered for each ORF. Additional complexity was related to the necessity to consider patterns related to regulatory sites. This task was possible to solve for each possible start separately, without getting into combinatorial complexity of possible combinations of starts over the whole contig. Technically, for a given gene start candidate, for each taxonomic group and genetic code we had to find the model cluster *h*, delivering the highest probability of the upstream sequence among possible clusters from cluster H. We had

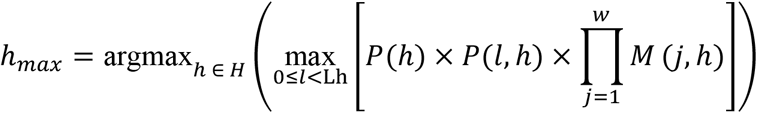

where P(*h*) was a prior probability of the cluster h from all possible clusters in H; P(*l, h*) was a probability of motif spacer with length *l* for the model cluster h; L_h_ was the maximum allowed length of a spacer; the value of 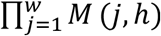 determined a likelihood of sequence with length *w*, the one ending at position –(*l*+1) from the putative gene start, to appear among sequences of modeled regulatory motifs.

The value of *h*_*max*_ delivered the probability

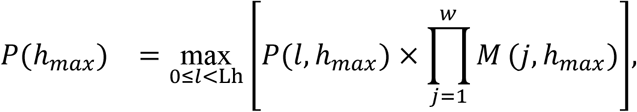

used in the Viterbi algorithm that was run on the given contig, with locally GC dependent models of each taxonomic group and genetic code. In each run the most likely parse for the given contig was determined along with its probability. These probability values were compared to identify the parse with highest probability among the candidate parses computed for each group Γ in the collections *G*_11_and *G*_4_.

Note that the probability computed in the Viterbi algorithm does not include prior probability *p*_*H*_(*h*). This exclusion is done to maintain the balance between the probabilities of sequences of motifs and probabilities of other sequence components (coding regions and start-contexts) used in the Viterbi algorithm. To account for a possibility that a contig comes from a genome with leaderless transcription, merged models of promoter and RBS were used for analysis.

### Gene prediction accuracy criteria and test sets

In case of complete genomes, errors in gene prediction may occur in gene starts while the gene ends could be found correctly. The opposite situation, an incorrect gene stop prediction when the start is predicted correctly normally does not occur. Empirically, incorrect prediction of position of the prokaryotic gene end (stop codon) and hence the gene reading frame by the current algorithms is happened rarely. Let assume that we have a test set of genes, designated as *T*. For a set of genes *P*, predicted in the same set of sequences, we define *M*_3_(*P, T*) the number of genes in *P* that have matching gene-ends in T, and |T| and |P| are numbers of genes in the sets T and P, respectively. Then, *whole gene* prediction Sensitivity (Sn) and Specificity (Sp) are:

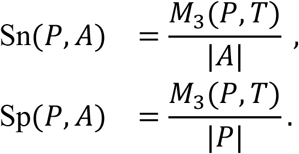

Then, let *M*_5_(*P, T*) designates a set of genes in *M*_3_(*P, T*) for which predicted starts match annotated starts. The error rate in start prediction is

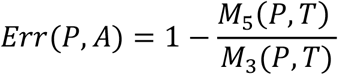

In case of metagenomic contigs the same types of errors would occur in gene prediction. To assess the accuracy of a *whole gene* prediction we used a set of short contigs generated by splitting the RefSeq genomes into pieces. To get the test set T_1_, a genome annotation defined by the PGAP pipeline (Tatusova et al. 2016; Haft et al. 2018) was mapped to contigs originated from the same genome.

For assessing the accuracy of gene start prediction, we had two sets, that we would designate as T_2_ and T_3_. The set T_2_ was the set of genes with experimentally verified starts. The set T_3_ was compiled as follows. In each genome annotated by the PGAP pipeline the gene starts were determined either with support of protein homology or if no such support was available, the gene start was determined by the *ab initio* prediction made by GeneMarkS-2. To eliminate influence of GeneMarkS-2 we consider only the group of genes with starts inferred from the protein alignment evidence. Further on, in this group we selected genes whose starts coincided with predictions of StartLink+ (Gemayel et al. 2021) supposed to have error rate ∼ 1%. This set was designated as T_3_.

### Motif models in genomes with reassignment of TGA

Among 101 prokaryotic genomes where TGA stop codon was reassigned to a codon for Trp (genetic code 4) 58 genomes were classified as group A, 33 as group C, and 5 each as groups B and X. The absence of A* and D* was to be expected since TGA in known archaeal species had always a role of a stop codon. The small sample sizes in groups B and X precluded further analysis. Interestingly, these bacterial genomes from groups A (RBS with the SD consensus) or group C (leaderless transcription) had low GC-content (Fig. 4).

**Figure 4.**
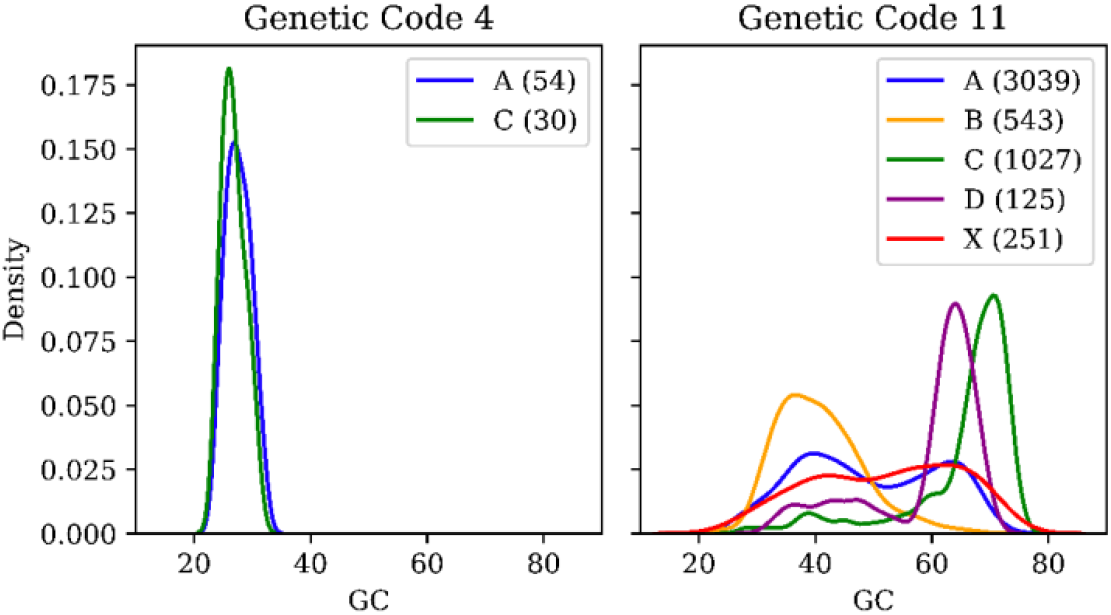
The per-group density distributions of representative bacterial genomes across GC, for genetic code 4 (left) and 11 (right). The number of genomes per group is shown in each panel.

In genomes where TGA was a stop codon (code 11), almost all genomes in groups C and D had high-GC content (Fig.4). In 33 genomes of group C with code 4, the percentage of genes with leaderless transcripts ranged from 30% to 60%, with an average of 46% (Fig. 5). This type of distribution of fraction of genes with leaderless transcription was similar the one observed in genomes of group C having code 11 (Lomsadze et al. 2018). Also, in genomes with code 4, e.g., in the four *Mycoplasma* genomes, group C, spacer length distributions for RBS motifs, as well as for promoter motifs had unimodal shapes (Fig. 6, bottom panel). These distributions were similar to the ones observed in genomes of group C with code 11 (Lomsadze et al. 2018). The motif models for the genomes of groups A and C having code 4 were included into the MetaGeneMark-2 model library.

**Figure 5.**
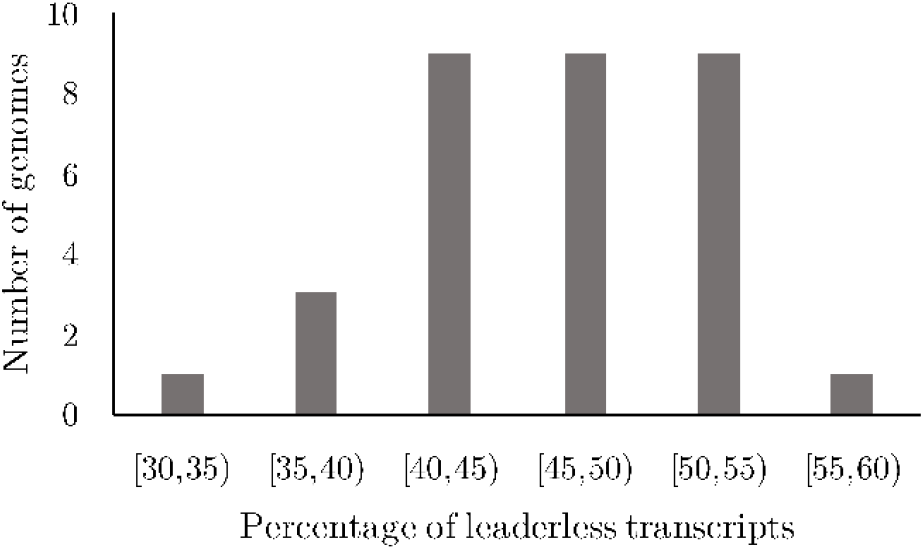
Distribution of the percentage of transcripts predicted as leaderless by GeneMarkS-2, in 33 genomes with genetic code 4 (the genomes from group C).

**Figure 6.**
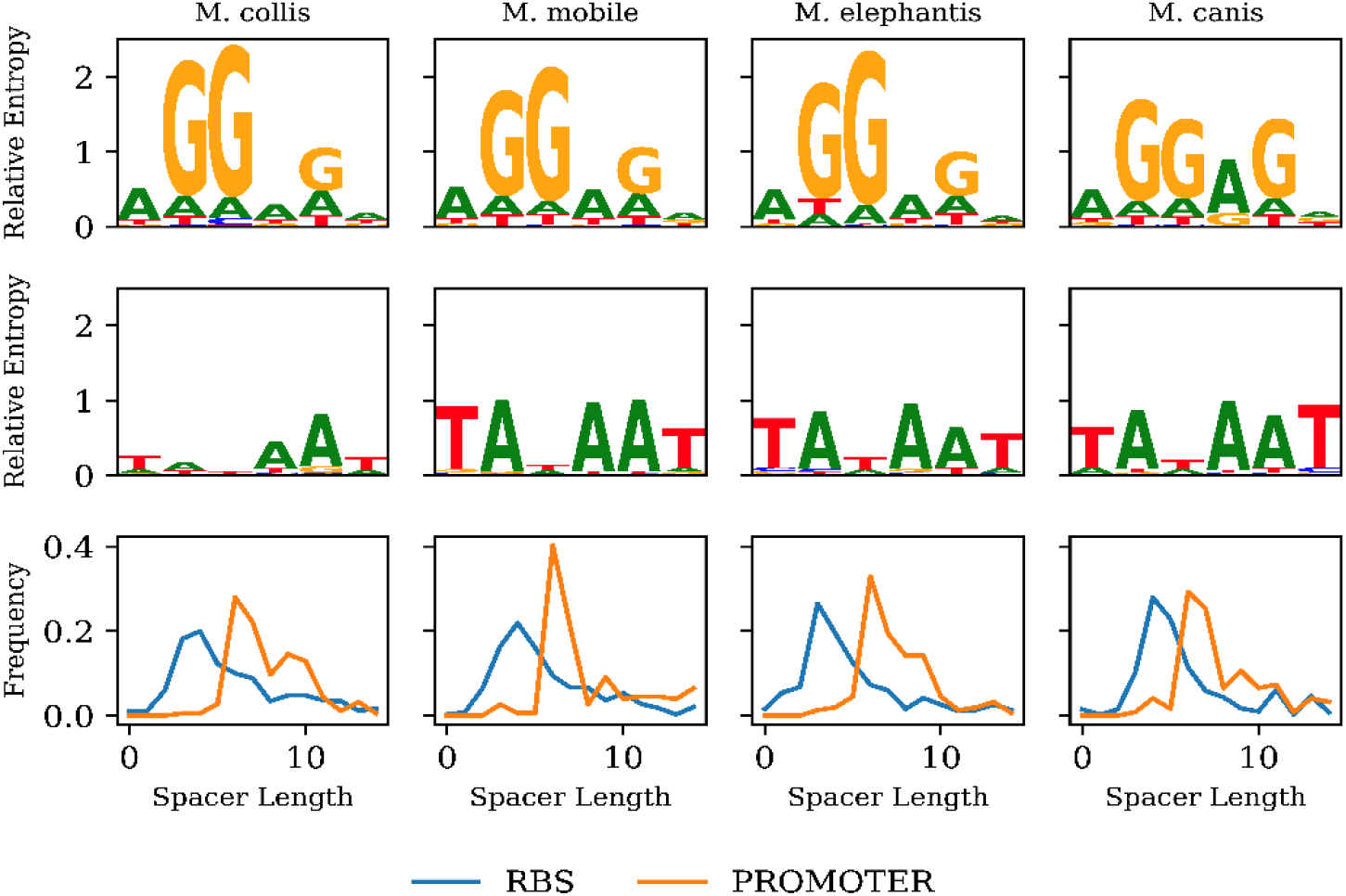
Relative entropy logos of the RBS models (top row) and the promoter models (middle row) derived for the four Mycoplasma genomes. The spacer length distributions are shown in the bottom row.

### Identification of the TGA role in metagenomic contigs

Identification of a role of TGA in a contig in MetaGeneMark-2 run was a part of model selection for genomic groups Γ from the collections *G*_11_and *G*_4_ (see 3.4). To assess the accuracy of the identification of the role of TGA, i.e., the genetic code 11 or 4, we used contigs of a simulated metagenome. For MetaGeneMark-2 and for MetaProdigal, we made the accuracy assessment on a simulated metagenome created from 100 genomes with genetic code 11 and from 5 genomes with genetic code 4, all genomes selected at random from 11,706 genomes annotated by NCBI (see Materials). Complete genomic sequences were split into contigs with length selected at random from the distribution of contigs length observed in assembled metagenome GCA_000763215 (GenBank). The simulated metagenome contained 67,200 contigs with total length 429 Mb, and minimal contig length 1,000 nt.

## Results

### Evaluation of the gene start prediction accuracy

Experiments aimed to experimentally verify gene translation starts had been conducted earlier for eight prokaryotic species (Table 1). These efforts resulted in translation start verification of 3,451 genes; all this data was used for assessing the error rates of MetaGeneMark-2 and of the four other metagenomic gene finders. To make an unbiased test of MetaGeneMark-2, we excluded the eight genomes (Table 1) from the sequence data used for the derivation of site models generating functions. The five gene prediction tools were run on each of the eight complete genomes; less than 1% of the genes with experimentally verified starts was missed by each tool. We observed (Table 1) that MetaGeneMark-2 made the least number of errors – 178 (5.16%), followed by MetaProdigal - 247 (7.16%) and MetaGeneMark – 477 (13.8%). We should mention that the MetaProdigal test on the *E. coli* genome was not entirely unbiased. The same set of the *E. coli* genes with verified starts was used in Prodigal (a precursor of MetaProdigal) for a discriminative training of the gene start model (Hyatt et al. 2010). However, MetaGeneMark-2 made less errors in predicting gene starts even in comparison with Prodigal, the whole genome gene finder (Table 1).

**Table 1.** Statistics of errors in gene start predictions for 3,451 genes with experimentally verified starts. Five metagenomic gene finders were compared in this test along with Prodigal and GeneMarkS-2 designed for single species genomic assemblies. Highlighted were the lowest rate per species (black, bold) and the second lowest rate (red, bold). All the gene finders were run on the full genomic assemblies with predictions recorded only for the verified genes

| Tool | Meta<br>GeneMark | Frag<br>GeneScan | MetaGene<br>Annotator | Meta<br>Prodigal | Meta<br>GeneMark-2 | Prodigal | GeneMarkS-2 |
| --- | --- | --- | --- | --- | --- | --- | --- |
| Species |  |  |  |  |  |  |  |
| *A. pernix* | 40 | 62 | 16 | **3** | 19 | **3** | 6 |
| *D. deserti* | 45 | 78 | 43 | 56 | **20** | 48 | **14** |
| *E. coli* | 129 | 101 | 34 | **19** | 34 | **18** | 28 |
| *H. salinarum* | 31 | 118 | 45 | 44 | **10** | 15 | **7** |
| *M. tuberculosis* | 134 | 117 | 97 | 86 | **67** | 77 | **65** |
| *N. pharaonis* | 13 | 52 | 19 | 6 | **4** | 5 | **3** |
| *R. denitrificans* | 74 | 50 | 43 | 25 | **17** | 26 | **19** |
| *S. PCC* | 11 | **4** | 12 | 8 | 7 | **4** | 5 |
| **Total** | 477 | 582 | 309 | 247 | **178** | 196 | **147** |

In the second test set for assessing accuracy of gene start prediction, [StartLink+], we used genes selected from 427 genomes (see Section 3.4). The lowest observed error rate – 6% was observed for MetaGeneMark-2, followed by MetaProdigal – 8%, MetaGeneAnnotator – 11%, MetaGeneMark – 13% and FragGeneScan – 14% (Fig. 7). It is worth to note that the error rates (in percentages) recorded in experiments with the [StartLink+] set were consistent with the error rates seen for the first test set. Similarly to the observations for the first test set, all the tools but FragGeneScan missed less than 1% of the genes in the [StartLink+] test set, while FragGeneScan missed 2.6% (Fig.8).

**Figure 7.**
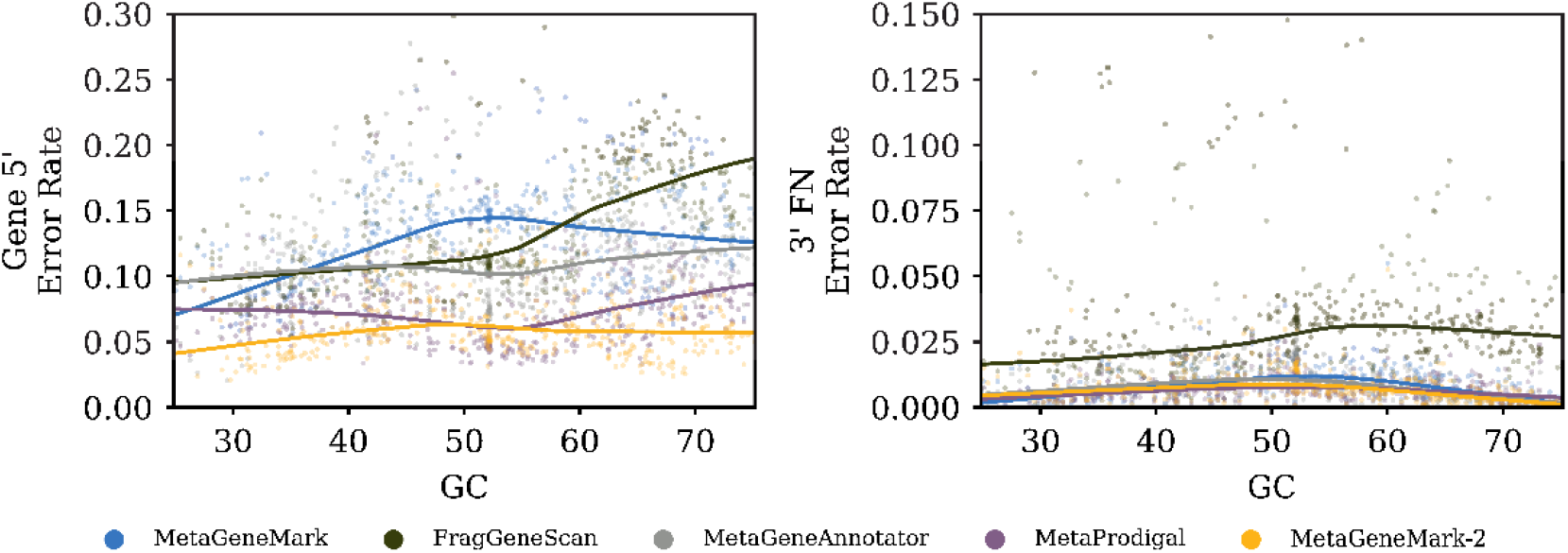
The error rates observed in predictions of gene starts (5’ ends) by the five gene finders shown as functions of genome GC content. The test set T_3_ used genes from 427 genomes. For genomes with GC content below 30% and higher than 70% we used models derived for genomes with 30% and 70% GC, respectively.

Out of the eight species, bacteria *M. tuberculosis, D. deserti*, and in archaea *H. salinarum* are known for frequent leaderless transcription. In genomes of these three species, we considered 142 verified genes whose starts have been predicted correctly by MetaGeneMark-2 but not MetaProdigal (Table 2a). Also, we considered 53 genes whose starts were predicted correctly by MetaProdigal but not MetaGeneMark-2 (Table 2b). We observed that 123 (86.7%) out of 142 gene starts were predicted by the MetaGeneMark-2 model designed for leaderless transcription (Table 2a). The gene upstream motifs derived by MetaGeneMark-2 for these 123 genes (Fig. 8) exhibit typical patterns for promoter sequences. The promoter spacer length distributions had an average length 6 *nt* in bacteria and 23 *nt* in archaea. Incidentally, for 83 (68%) of these 123 genes MetaProdigal was incorrectly using the RBS type model. For the 53 genes predicted correctly by MetaProdigal with the RBS model (Table 2b) but not by MetaGeneMark-2, the later was using the RBS and non-RBS models at about equal rate.

**Table 2.** (**A**) Statistics of models used for genes with verified starts in *D. deserti, M. tuberculosis* and *H. salinarum* for which MetaGeneMark-2 made correct gene start prediction but not MetaProdigal. Shown are the MetaGeneMark-2 model types chosen for these genes (RBS or leaderless), and the MetaProdigal model types (RBS or No RBS). (**B**): Same as in panel a/ for genes for which MetaProdigal made correct predictions, while MetaGeneMark-2 did not.

| MetaGeneMark-2 matching verified starts |  | MetaProdigal |  |
| --- | --- | --- | --- |
|  |  | RBS | No RBS |
| RBS | 19 | 16 | 3 |
| Leaderless | 123 | 83 | 40 |

**B**
| MetaProdigal matching verified starts |  | MetaGeneMark-2 |  |
| --- | --- | --- | --- |
|  |  | RBS | Leaderless |
| RBS | 20 | 16 | 4 |
| No RBS | 33 | 17 | 16 |

**Figure 8.**
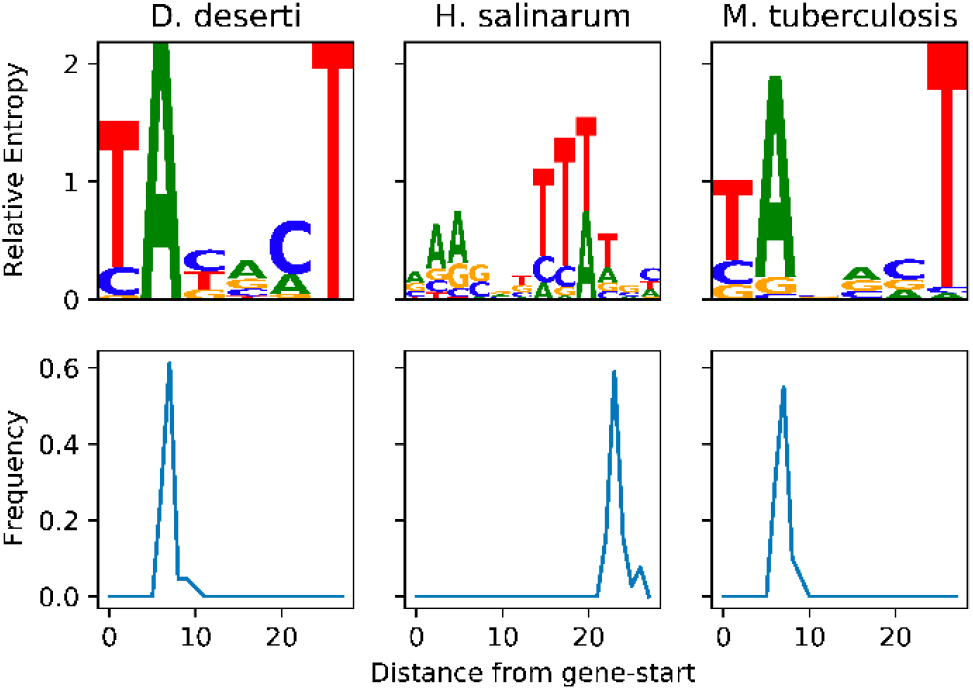
The relative entropy logos of the promoter motifs (top panel) along with the spacer length distributions (bottom panel) shown for 123 genes (44 from *D. deserti*, 39 from *H. salinarum* and 40 from *M. tuberculosis*) which MetaGeneMark-2 predicted as genes with leaderless transcription. Positions of the predicted starts of the 123 genes matched the experimentally verified positions (Table 1).

### Gene-level accuracy determined for complete genomes

To determine the accuracy in prediction of gene 3’ ends we made another test of the five gene finders run on 427 complete genomes (Fig. 9). We must note that, unlike the four other tools, MetaGeneAnnotator was designed to change its mode of operation when running on contigs longer than 5,000 nt. On such contigs, MetaGeneAnnotator performed a whole cycle of self-training. Thus, in a test on a complete genome MetaGeneAnnotator operated as a whole genome gene finder rather than a metagenomic one. Therefore, the accuracy of MetaGeneAnnotator we could observe in such an experiment presents an upper limit for a metagenomic gene finder.

**Figure 9.**
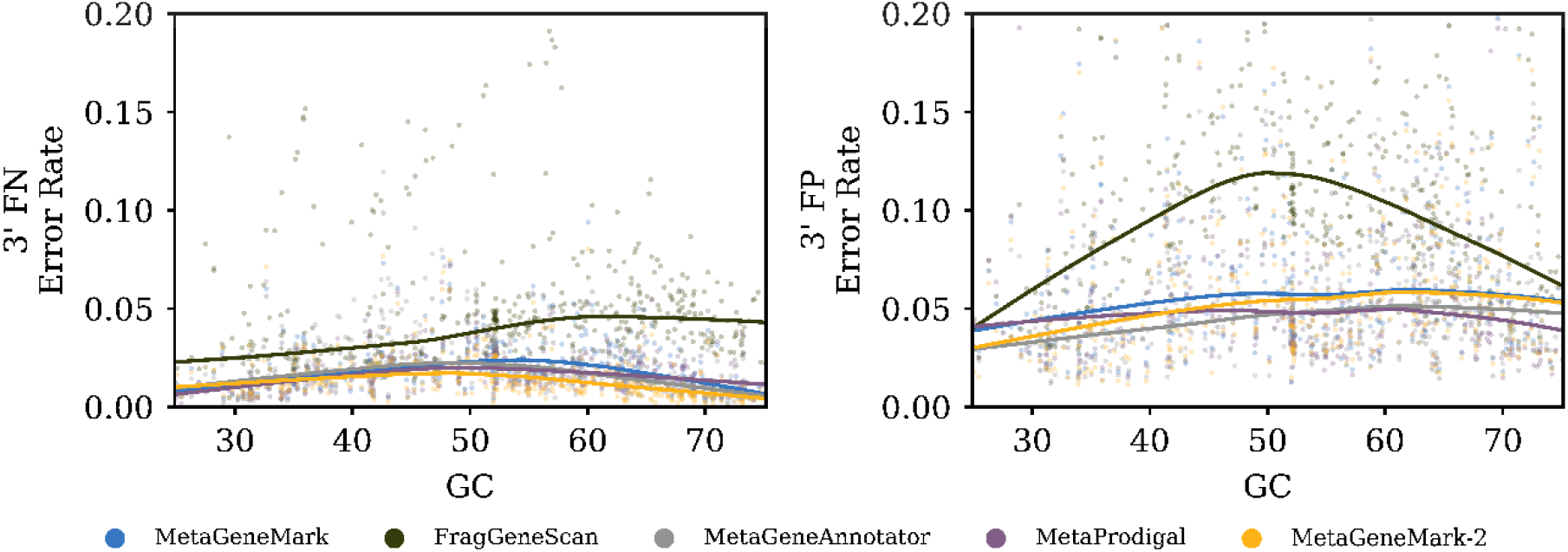
The false negative and false positive error rates of the whole gene prediction (misplacing the gene 3’ end) by the five gene finders. The error rates were computed by comparison with the RefSeq annotation of 427 complete genomes. The GC content of the 427 genomes varied from 25% to 75%.

### Gene prediction in simulated metagenomic contigs

We used the same 427 genomes to generate sets of contigs having equal length, 250, 500, 1000 and 1500 nt. Starts of the genes predicted in each contig were compared with annotations of gene starts of test set T_3_ (see 3.5) and transferred from the corresponding genomes (Fig. 10). MetaGeneMark-2 showed the lowest error rate, uniformly with respect to contig GC, in the contigs shorter than 1000 nt. For the longer fragments MetaGeneMark-2 had the lowest error rate for GC content lower than 45% and higher than 60%. In the interval from 45% to 60% GC content it had slightly higher error rate than MetaProdigal (Fig. 7).

**Figure 10.**
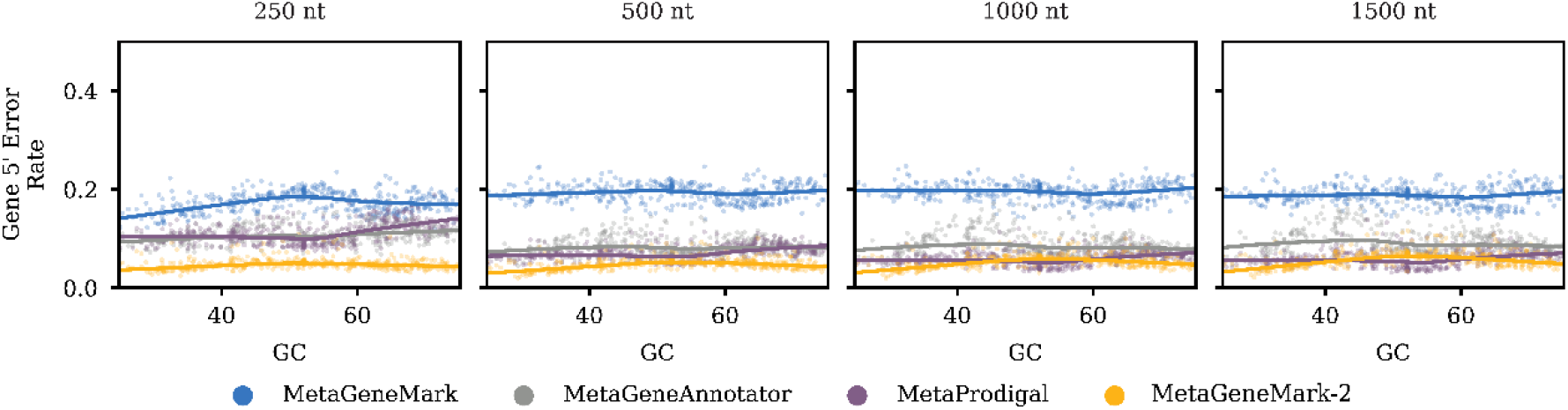
The gene start prediction error rate observed in equal length contigs from 427 genomes. The starts of the genes were compared with starts determined in the test set T_3_.

We also compared predictions of the gene 3’ ends made in the simulated metagenomes by the five gene finders with the annotation of genes in the set T_3_. For contigs with lengths from 250 to 1,500 nt we observed average false negative and false positive rates to be below 2% (Fig. 9), consistent with the benchmarking on complete genomes.

### Identification of a role of the TGA codon

Here we compared performance of MetaGeneMark-2 and MetaProdigal, the only two computational tools making recognition of the TGA role in an anonymous contig. MetaGeneMark-2 correctly identified a role of TGA in all the test genomes (185 genomes with genetic code 4 and 11,521 genomes with genetic code 11). MetaProdigal identified TGA as a stop codon in all 18 *Hodgkinia* genomes where TGA codes for Trp (code 4) and incorrectly reassigned TGA as a sense codon in 51 genomes where TGA was a stop codon (code 11).

The accuracy of gene 3’ end predictions could be affected significantly when a role of TGA is assigned incorrectly. To illustrate this point, we selected 5 genomes where the role of TGA was identified differently by MetaGeneMark-2 and MetaProdigal. For subsets of genes supported by alignments to homologous proteins (Haft et al. 2018), gene predictions made by each gene finder were compared with the RefSeq annotations. The gene prediction error rate of MetaProdigal was 20.4%, while MetaGeneMark-2 it was 0.7% (Table 3). Notably, the whole genome gene finder, Prodigal, (Hyatt et al. 2010) with a model correctly reflecting the TGA role showed ∼0.8% difference with the RefSeq annotation.

**Table 3.** Error rate in predictions of 3’ end of genes by MetaGeneMark-2 and MetaProdigal in five genomes where the two gene finders identified the role of TGA differently. The test set contained only those genes whose RefSeq annotations were supported by alignments of homologous proteins.

| Species | # of PGAP genes supported by homology | % of genes missed by MetaGeneMark-2 | % of genes missed by MetaProdigal |
| --- | --- | --- | --- |
| *Acholeplasma axanthum strain NCTC10138* | 1,389 | 1.2 | 17.1 |
| *Anaerococcus obesiensis ph10* | 1,914 | 4.0 | 18.5 |
| *Helicobacter bizzozeronii CIII-1* | 1,613 | 6.7 | 32.9 |
| *Peptoniphilus duerdenii ATCC BAA-1640* | 1,789 | 2.2 | 23.5 |
| *Sneathia amnii strain SN35* | 1,084 | 1.7 | 19.2 |

Finally, when MetaProdigal and MetaGeneMark-2 were run on the simulated metagenome with length 429 Mb (see 3.6.1), we observed that the role of TGA codon was incorrectly assigned by MetaProdigal and by MetaGeneMark-2 in 1.07% and 0.17% of contigs, respectively.

### Merging sequence motif models

In the GeneMarkS-2 algorithm RBS sequences were represented by positional frequency models with 24 parameters. We visualized the distribution of distance between the vectors of the RBS models for 800 archaeal and 5,200 bacterial genomes by the UMAP package (McInnes et al. 2018) that mapped these vectors from the 24D space into a 2D space. After the mapping, the 2D points were colored to depict 1/ the species domain, i.e., archaea or bacteria, 2/ the genome group, 3/ the genome GC content (Fig. 11).

**Figure 11.**
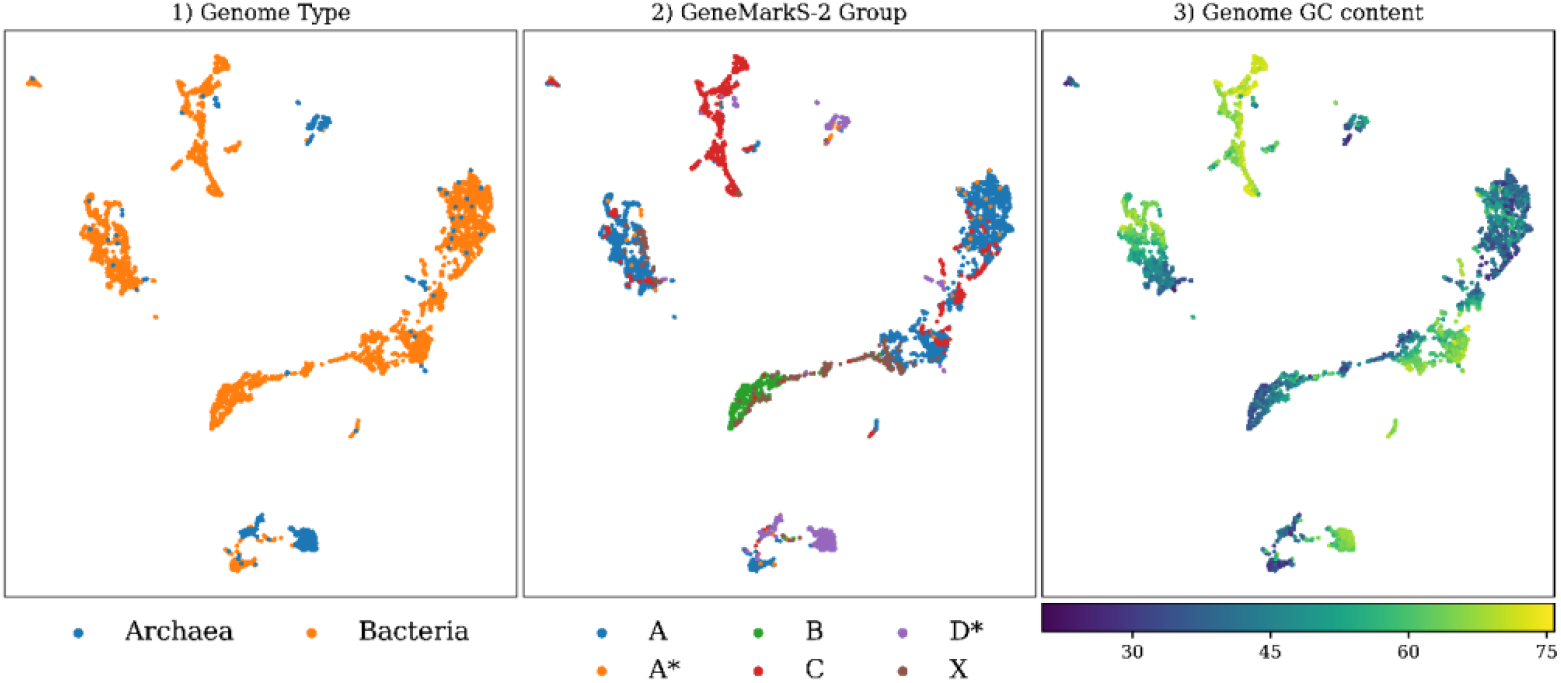
A 2D graph depicting clustered projections of the RBS models with colors corresponding to 1/ archaea and bacteria; 2/ six genome groups; 3/ different GC content.

The mapping shows separation of the models built for archaeal genomes from those of bacterial genomes. There is also a clear separation between models related to different GeneMarkS-2 genome groups (Lomsadze et al, 2018). Interestingly, Group C cluster correlates with the cluster of the models for high GC genomes. The AT-rich non-SD RBS models for the Group B bacterial genomes appear in the AT rich genomes. Also, the models for Group C genomes are separated from the models for Group A, despite both having the SD RBS.

Overall, this analysis provides strong supporting arguments for inferring parameters of models the gene upstream regulatory regions with help of generating functions dependent on the sequence GC content and the taxonomic origin.

### Running time

A run of MetaGeneMark-2 on the 134 Mb sediment metagenome did take 24 minutes (www.ncbi.nlm.nih.gov/search/all/?term=GCA_000763215); a MetaProdigal run on the same metagenome was finished in 34 minutes.

## Discussion

Automatic computational analysis of volumes of new metagenomic sequences is a critically important element of the high throughput biotechnology created to penetrate mysterious depths of the astronomically vast world of microbes. Metagenomic gene finders make a first cut in the process of reconstruction of the protein complements and biological networks in microbial communities. Improving accuracy of state-of-the-art gene finders is not a trivial task. Focus of this work was on improving gene start prediction. The task of recognition of a short sequence containing gene start had to be solved for an anonymous sequence. The difficulty of modeling such sequences was aggravated by the possible presence of multiple types of regulatory sites – SD-RBS, non-SD RBS, bacterial promoter, archaeal promoter, etc. Therefore, the method of *parameters generating functions* introduced earlier (Besemer and Borodovsky 1999; Zhu et al. 2010) was necessary to generalize beyond the initial boundaries. The new method had to automatically generate a model structure along with selection of parameter generating function, or even find the best model structure in the process of analysis.

Our task was facilitated by recent research (Lomsadze et al. 2018) that introduced a method of classification of regulatory sites of prokaryotic genomes and, subsequently, more detailed modeling of sequences near translation initiation sites. The main types of models considered here were Shine-Dalgarno and non-Shine-Dalgarno ribosome binding sites as well as promoter sites situated, in case of leaderless transcription, close to gene starts.

Expanding a very brief outline given above, we should say that yet for the original MetaGeneMark (Zhu et al. 2010) we demonstrated that the codon frequencies and, more generally, the k-mer frequencies in protein-coding regions could be sufficiently accurately represented by non-linear functions of genomic GC content. These GC dependent *parameters generating functions*, were used in MetaGeneMark to determine parameters of models of protein-coding regions for each metagenomic contig. A similar approach was used for generating parameters of the models of non-coding regions. Notably, the position specific frequency matrices for ribosome binding sites and promoters together with the spacer length distributions along a few sequence positions, had smaller number of parameters than models of protein-coding regions. Still, the simpler structure of the additional models for MetaGeneMark-2, was ‘compensated’ by the larger number of competing models existing in the same range of GC composition. It should be fair to say that in MetaGeneMark a competition of models within the same GC bin was also present. It was a competition of just two protein-coding region models derived for bacterial and archaeal domains.

Modeling of sequences near translation initiation site was used in other gene finders. MetaProdgal has 50 models of protein-coding regions complemented by the species-specific models of RBS and non-RBS starts. The 50 genomes with various GC content were selected from 1,415 genomes to reach best average performance on the whole set of genomes (Hyatt et al. 2012). We observed that for contigs with GC content in the mid GC range accuracy of MetaProdigal and MetaGeneMark-2, was about the same (Fig. 7). However, in low and high GC ranges, with more contigs coming from genomes with leaderless transcription, advantage of MetaGeneMark-2 became significant (Fig. 7).

Genomes of species known for utilizing mechanisms of leaderless transcription, *D. deserti, H. salinarum* and *M. tuberculosis*, contained a total of 1,615 genes with experimentally verified starts; on this test set MetaGeneMark-2 made 89 fewer gene start prediction errors than MetaProdigal (Table 1).

We saw that translation initiation site model of MetaGeneMark-2 was almost as informative & efficient as the model used in GeneMarkS-2; gene start prediction accuracy by MetaGeneMark-2 is close to one of GeneMarkS-2. Switching off the translation initiation site model led to immediate degradation of the accuracy of the gene start prediction. Better performance of MetaGeneMark-2 could be explained by the use of significantly more diverse set of the translation initiation site models. Particularly, MetaGeneMark-2 employed five types of translation initiation site models (for genome groups A,B,C,D and X as described in (Lomsadze et al. 2018)). Parameters sets of these models were pre-computed in GC range from 30% to 75% GC (a total of 112 models: 80 with code 11 and 32 with code 4).

We observed that most of the parameters in a model of protein-coding region remained the same regardless of the role of TGA. Differences appeared only in the parameters related to frequencies of k-mers containing the TGA triplet.

Among 50 species-specific models of MetaProdigal, there were four for the *Mycoplasma* species where TGA codon codes for Trp. MetaProdigal identified TGA role correctly in all the *Mycoplasma* contigs which GC content was in the 22-40% range. However, in genomes of 18 *Candidatus Hodgkinia cicadicola*, with GC content in the 38-58% range, the role of TGA was misidentified. Another type of error, predicting TGA, being a stop codon, coding for Trp, was made by MetaProdigal in 15 genomes where the spectrum of hexamer frequencies in protein-coding regions was similar to the spectrums observed in the *Mycoplasma* genomes (Table 3 shows results for 5 out of 15 genomes). MetaGeneMark-2 made lower number of errors in identification of a role of TGA; it showed consistently good accuracy for prokaryotic species across a wide range of genomic GC content.

All over, we demonstrated that in multiple tests a new metagenomic gene finder, MetaGeneMark-2, delivered better accuracy of gene start prediction in metagenomic contigs than several state-of-the-art computational tools.

## Availability of the code and datasets

The MetaGeneMark-2 software is available for academic users at https://github.com/gatech-genemark/MetaGeneMark-2. All scripts and data used to generate figures and tables for this manuscript are available at *https://github.com/gatech-genemark/MetaGeneMark-2-exp*.

## Author Contributions

Conceptualization, M.B., A.L. and K.G.; methodology, K.G., A.L. and M.B.; software, K.G., A.L.; writing - original draft preparation, K.G. and M.B.; writing - review and editing, M.B.; visualization, K.G.; funding acquisition, M.B. All authors have read and agreed to the published version of the manuscript.

